# Linking indices for biodiversity monitoring to extinction risk theory

**DOI:** 10.1101/000760

**Authors:** Michael A. McCarthy, Alana L. Moore, Jochen Krauss, John W. Morgan, Christopher F. Clements

## Abstract

Biodiversity indices often combine data from different species when used in monitoring programs. Heuristic properties can suggest preferred indices, but we lack objective ways to discriminate between indices with similar heuristics. Biodiversity indices can be evaluated by determining how well they reflect management objectives that a monitoring program aims to support. For example, the Convention on Biological Diversity requires reporting about extinction rates, so simple indices that reflect extinction risk would be valuable. Here we develop three biodiversity indices that are based on simple models of population viability that relate extinction risk to abundance. The first index is based on the geometric mean abundance of species. A second uses a more general power mean. A third integrates both the geometric mean abundance and trend. These indices require the same data as previous indices, but they also relate directly to extinction risk. Field data for butterflies and woodland plants, and experimental studies of protozoan communities show that the indices correlate with local extinction rates. Applying the index based on the geometric mean to global data on changes in avian abundance suggests that the average extinction probability of birds has increased approximately 1% from 1970 to 2009.

## INTRODUCTION

The importance of biodiversity for a healthy and equitable society has been acknowledged by over 190 countries that ratified the Convention on Biological Diversity (CBD). The convention has a specific target to reduce the extinction risk of species (Secretariat of the Convention on Biological Diversity 2010), so monitoring of species extinction is important. Reporting actual extinctions, while potentially informative, is retrospective, whereas the convention and many other biodiversity programs seek to reduce future extinctions. Further, retrospective assessments are subject to error because the fate of species is known imprecisely (Collar 1998; Keith & Burgman 2004; Rout et al. 2010). Hence, biodiversity monitoring programs would be more valuable if they can be interpreted in terms of extinction risk.

Changes in the assessed risk to species can contribute to biodiversity monitoring. For example, the IUCN Red List is used to calculate the Red List Index (RLI, Butchart et al. 2007), one of four global indicators of biodiversity status and trends approved by the CBD (Jones et al. 2011). The relationships of the other three indicators (extent of forest; protected-area coverage; and the Living Planet Index, LPI, Jones et al. 2011) to extinction risk are not explicit.

Buckland et al. (2005) identified three aspects of species diversity that are of primary interest when monitoring changes over time: number of species, overall abundance and species evenness, from which they derived six desirable criteria for an index of biodiversity based on abundance data. On evaluating several proposed indices against these criteria, the geometric mean of relative abundances was one of only two that met all six criteria, with van Strien et al. (2012) lending further support to the geometric mean.

While we agree with the heuristic properties used to assess different indices of biodiversity, a good index should also be clearly related to particular management objectives or biodiversity outcomes. For example, where extinction risk is the management concern, understanding how the index reflects changes in this risk would be desirable. In the absence of a single measurable definition of biodiversity (Secretariat of the Convention on Biological Diversity 2010; Jones et al. 2011), we aim to examine how abundance data might be used to monitor extinction rates of species for the purposes of reporting under the CBD and other biodiversity programs.

Here, we use simple models of population viability to develop three indices of extinction risk based on abundance data. These indices are designed to have the same data requirements as those considered by Buckland et al. (2005), but with the additional benefit of being directly related to extinction risk. We evaluate the indices using simulation, field data on local extinctions of butterflies and woodland plants, and experimental data on protozoan communities. Finally, we interpret changes in the LPI in terms of changes in the average probability of extinction of species.

## Methods

The indices are derived from simple models of population viability, using clearly articulated assumptions that can be tested. First, consider the case when the long-term average population growth of each species is negative. If we assume that each species is experiencing deterministic exponential decline, then

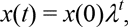

where *x*(*t*) is population abundance at time *t,* and *λ* is the growth parameter (*λ* < 1 for a declining population). It is then straightforward to calculate that extinction (such that *x*(*t*) = 1) occurs at time *T* =–ln[*x*(0)]*/* ln[*λ*]. If the long-run growth rate is negative, then for stochastic population models the mean extinction time is also approximately logarithmically dependent on initial population size (Lande 1993).

With the simplifying assumption that the rate of decline is the same for each species (we address this particular assumption later), the mean expected time to extinction, averaging over *n* species, is proportional to the mean of the logarithm of population abundance. As we show below, the mean expected time to extinction is proportional to the logarithm of the geometric mean of population abundances (*M*_0_);

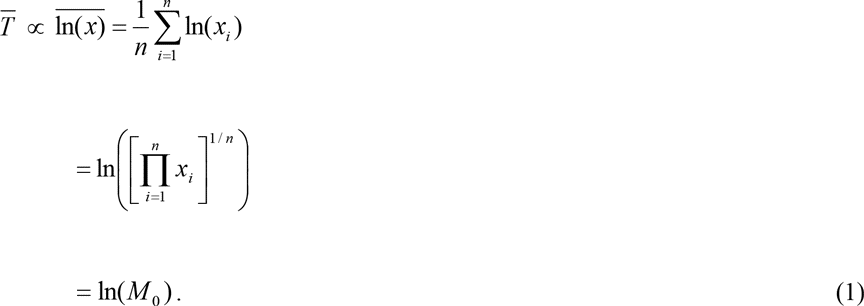

Equation 1 relates the mean time to extinction to the geometric mean abundance. However, it would be helpful to determine how this index might relate to the proportion of species going extinct. We approximate this by assuming that times to extinction have an exponential distribution. The proportion of species going extinct within time *t* is then 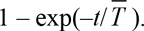. When this proportion is ≤0.2, it can be approximated by 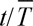, leading to:

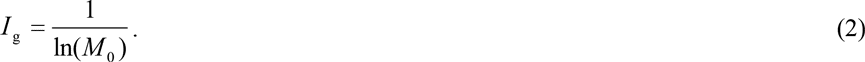

This index should correlate linearly with the proportion of species going extinct under the assumptions stated above. The approximation of 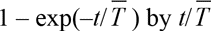 will tend to lead to non-linearity (but a monotonic relationship) for higher risks.

We develop a second index based on a different set of assumptions. We consider a stochastic population model in which the logarithm of the population growth rate has a normal distribution with a mean of zero and variance *σ*^2^. For this model, the risk of extinction within a given time period *t* is (Ginzburg et al. 1982; Dennis et al. 1991; McCarthy & Thompson 2001):

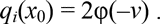

where φ() is the standard normal cumulative distribution function, *v* =–ln(1/*x*_0_)/(*σ*√*t*) and *x*_0_ is the initial population size. This functional form could be used as an index, but it does not provide a simple numerical solution. Instead, we approximated this equation by a function of the form *A x*^−*B*/(σ√*t*)^ (by approximating log(*q_i_*(*x*_0_)) as a linear function of log(*x*)) with the values of *A* and *B* depending on the value of the extinction risk. For small extinction risks, *q_i_<*≈ 0.15, *A* = 2.2 and *B* = 1.87 provide a good approximation. When the extinction risk is close to one, a better approximation is *A* =1 and *B* = 0.798. Regardless, the probability of extinction scales approximately with abundance in proportion to *x*^−*b*^, with *b* = *B*/(*σ*√*t*). Thus, averaged across *n* species, we would expect the proportion of species going extinct to be

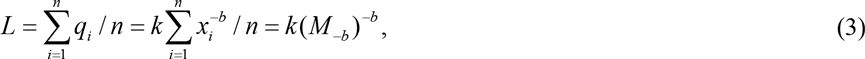

where *k* is a constant of proportionality and *M*_–*b*_ is a power mean of abundance with power *p* =–*b*,

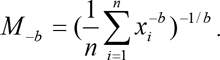

Consequently, our second index is based on a power mean of abundance:

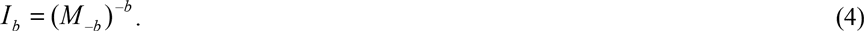

The value of *b* depends on the time horizon over which risks are assessed. If we consider a time horizon of *T* = 100 years and a standard deviation of *σ*= 0.1 (Dennis et al. 1991), the extinction risk of each species is likely to be relatively small (recall, zero mean growth rate), and *b* would be of the order 1.8 ≈ 2. The value of *b* will be larger for shorter time horizons.

A third index can be derived from the deterministic model that accounts for the population growth rate, in addition to population size. Noting again that the mean time to extinction under deterministic decline is–ln[*x*(0)]*/*ln[*λ*], then the proportion of species going extinct can be approximated by–ln[*λ*]*/*ln(*M*_0_), allowing communities with different population growth rates of species to be compared. Using the mean of the logarithmic population growth rate of species within a community, *μ_r_*, as the estimate of ln[*λ*] leads to the index:

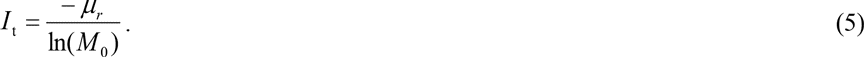

This index requires extra data, being the population growth rates of species within the community. Such data might be uncommonly available, but are necessary to compare risks among communities where the species are declining at different rates.

## Simulations for evaluating indices

We simulated stochastic species dynamics within communities to evaluate the correlation between the different indices and the proportion of species going extinct. Each community consisted of 500 species, and there were 100 different communities. For each species *j* in community *i*, we simulated the population dynamics over 20 time steps using the exponential growth model such that the population size in time *t* + 1 is given by:

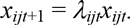

Parameter values for the 100 different communities were chosen such that the proportion of species going extinct spanned a wide range (in our case between 0.02 and 0.64). Within each community, the initial population size ln*x_ij_*_0_ was drawn from a lognormal distribution with mean *μ_N_* and coefficient of variation *c_N_*, and the logarithmic growth rate ln*λ_ijt_* was drawn from a normal distribution with mean *μ_r_* and standard deviation *σ_r_*. The proportion of 500 species that fell to or below one individual measured the average extinction risk of the community.

To ensure that each community had different initial population sizes and different trends in abundance (and hence different average extinction risks), the mean and coefficient of variation of the population size (*μ_N_* and *c_N_*) and the mean and standard deviation of population growth rate (*μ_r_* and *σ_r_*) of each was varied among communities. The coefficient of variation *c_N_* was drawn from a uniform distribution on the interval [0.5, 3.0]. The mean population size was equal to 1.2*^d^*100, where *d* was drawn from a uniform distribution on the interval [0, 20], so mean population size varied among communities over the interval [100, 3834]. The mean population growth rate (*μ_r_*) was drawn from a uniform distribution on the interval [–0.3,–0.1], and the standard deviation (*σ_r_*) was drawn from a uniform distribution on the interval [0.05, 0.4]. To test how differences in abundance, rather than population trend, influence the performance of the indices, data were also simulated with *μ_r_* set to–0.2 for all communities.

The three indices of extinction risk (*I*_g_, *I*_b_ and *I*_t_) were calculated for the simulated communities and the correlations between these and the proportion of species going extinct was examined. The performance of the arithmetic mean abundance and the modified Shannon diversity index of Buckland et al. (2005), other putative biodiversity measures, were also examined for the simulated data. For these two cases, we multiplied the indices by −1 so that the indices would be expected to be positively correlated with extinction risk.

## Data for evaluating indices

The correlation between the indices and local extinction risk was evaluated using field data on Lepidopetera (Krauss et al. 2003) and woodland plants (Sutton & Morgan 2009). Because data on population trends were unavailable for these datasets, only eqns 2 and 4 were evaluated. We evaluated all three indices with data from experimental protozoan communities (Clements et al. 2013). The original publications detail the data and its collection; some information is provided here for context (see also Supporting Information). The data sets we examined reported both extinctions of multiple species and information on initial abundances.

Each dataset included information on the abundance of each of the species in replicate local communities at a particular point in time, and data on the proportion of those species that went extinction over a subsequent period of time. For the protozoan community, estimates of abundance were available at multiple points in time prior to the period over which extinctions were assessed. For each dataset, we calculated the indices using the abundance data (and the trend data in the case of the index *I*_t_ for the protozoan dataset).

For each dataset, we calculated the correlation (with 95% confidence interval based on a *z*-transformation; Sokal & Rohlf 1981) between the value of each index and the proportion of species in each community going extinct. We also determined, via simulation, the correlations that would be expected if each index were perfectly correlated with extinction of species, while accounting for the finite number of species in each community (Supporting Information). This allowed us to determine whether the observer correlations were substantially different from what would be expected given the limitations of the datasets.

## Relating *I*_g_ to the Living Planet Index

The LPI is the geometric mean abundance of vertebrate species in a particular year divided by the geometric mean in 1970 (Loh et al. 2005; Collen et al. 2009). Therefore, the index based on the geometric mean can be related to the LPI simply as *I*_g_ = 1/ln(*c* LPI), where *c* is the geometric mean abundance in 1970. If *I*_g_ is proportional to the probability of extinction, as assumed in its derivation, LPI values can be converted to proportional changes in the probability of extinction of species, which will equal–ln(LPI) / [ln(*c*) + ln(LPI)]. We calculated this quantity for the world’s birds based on published avian LPI values (Baille et al. 2010).

These proportional changes depend on *c*, which is not well known. The arithmetic mean abundance of birds is thought to be approximately 10 million individuals per species but, because species abundance distributions are heavily right-skewed, the geometric mean will be substantially less (Gaston & Blackburn 2003). We estimated the global species abundance distribution of birds, and hence the geometric mean, by fitting a log-normal distribution to data on reported population size for the global list of 1253 threatened species on BirdLife International’s website (http://www.birdlife.org/datazone/species/search; accessed 20 December 2011) and assuming an arithmetic mean of 10 million birds per species. We assumed that abundances of the remaining 8663 non-threatened species were greater than 1000. In this case, and in cases where the data on threatened species were provided as ranges, we fitted the model assuming censored data. When an upper limit was not provided, we set the upper limit of 10 billion individuals for each species, which is greater than the reported abundance of passenger pigeons, the world’s most abundant bird prior to its extinction. The geometric mean of the resulting log-normal probability distribution was then calculated. The sensitivity of the results to the calculated value of *c* was examine by varying *c* by one order of magnitude and re-calculating the proportional changes in the probability of extinction.

## Results

For the simulated communities with variation in mean growth rate among communities, the index based on the power mean (*I*_b_) and the index based on the geometric mean (*I*_g_) were positively correlated with the proportion of species going extinct (Pearson product moment correlations *r* = 0.39 and *r* = 0.50, respectively). Spearman rank correlations were similar (*r*_S_ = 0.34 and 0.49 respectively). Variation in mean growth rates among communities explained much of the imperfect correlations; correlations for the index that is based on population trend were high (*r* = 0.96; *r*_S_ = 0.99 for *I*_t_), and were similarly high for the geometric mean index (*I*_g_) when all communities had the same mean rate of decline (*r* = 0.97 when *μ_r_* =–0.2 for all communities).

The index based on the geometric mean (*I*_g_) and the index that considers population trend (*I*_t_) were more strongly correlated with the proportion of species going extinct than either index based on the arithmetic mean or the Shannon diversity (*r* = 0.44 in both cases when *μ_r,i_* varied on the interval [–0.3,–0.1], and *r* = 0.94 and 0.91 respectively when *μ_r,i_* was–0.2 for all communities). The index based on the power mean (*I*_b_) was the least strongly correlated with the proportion of species going extinct (*r* = 0.39 when the mean population growth rate varied among communities, *r* = 0.66 when it was consistent); this might be expected given the strong influence of the population trend on the simulated extinction risks, whereas the index *I*_b_ assumed no trend. Note, the derivation of *I*_g_ included a trend, but it dropped out of the calculation of the index as a proportionality constant by assuming the same trend for all communities.

For the real communities, the index based on the geometric mean abundance (*I*_g_) and the index based on the power mean (*I*_b_) were positively correlated with the proportion of Lepidopetera and woodland plant species that went extinct (Fig. 1a,c; Fig. 2). The 95% confidence intervals for these correlation coefficients did not encompass zero. In contrast, the correlations for these indices were negative for the protozoan dataset (Fig. 1e,f), although the correlation for the index that included population trends was positive (*r* = 0.33; Fig. 1 g, Fig. 2). In this dataset, abundances were similar for most communities, so the indices spanned a narrow range of values. The 95% confidence intervals for the correlation coefficient were wide (Fig. 2), so the strength of the relationship could not be determined reliably for the protozoan dataset.

**Figure 1.**
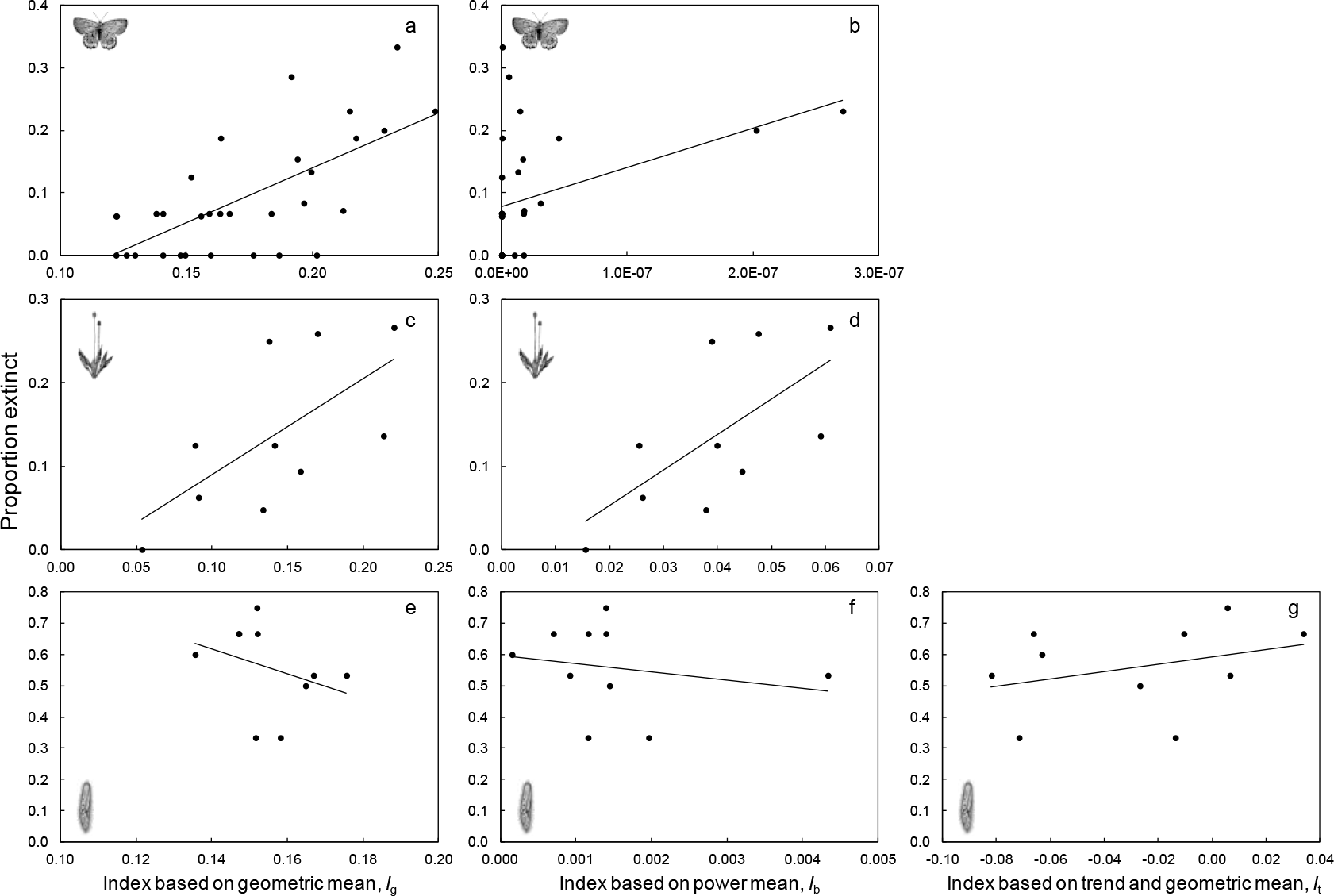
Relationship between the three different indices (*I_g_*, *I_b_*, *I_t_*) and the proportion of species going locally extinct for the three case studies: (a-b) for Lepidopetera; (c-d) for woodland plants; and (e-g) for protozoan communities. Each point represents a patch for the field studies (Lepidopetera and woodland plants) or the average of each type of community for the protozoan. The lines are linear regressions. Correlation coefficients with 95% confidence intervals are given in Fig. 2.

**Figure 2.**
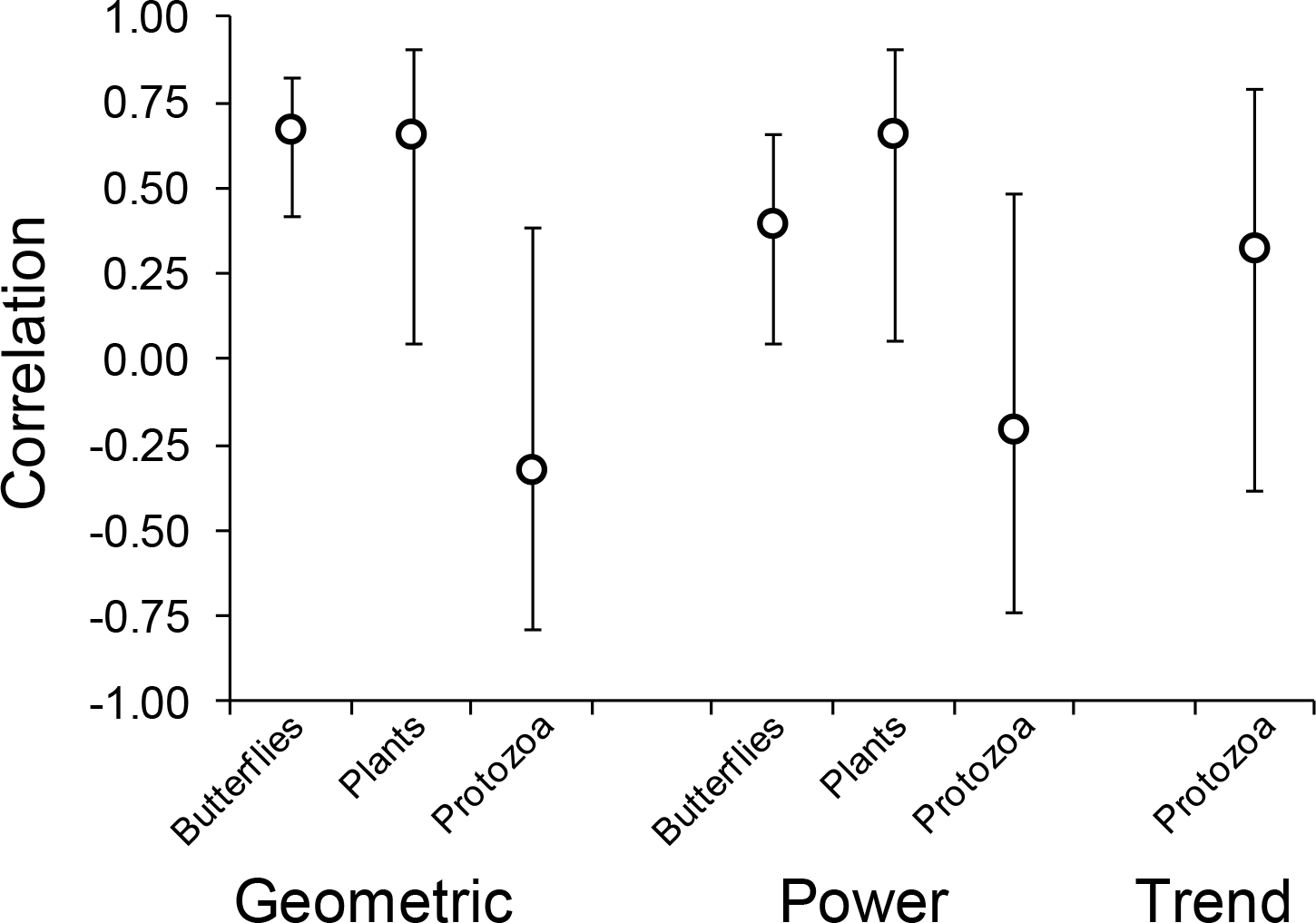
Observed correlation between the three indices (*I*_g_ based on the geometric mean; *I*_b_ based on the power mean; and *I*_t_ based on the geometric mean and trend) and the proportion of species going extinct from a community. Results are shown for each of the three different datasets (butterflies, plants, protozoa). The circles are the observed correlation coefficients and the bars are 95% confidence intervals.

There was only one case (the index based on the power mean for the protozoan dataset) that the observed correlation coefficient was both not significantly different from zero (Fig. 2) and substantially less than the correlation coefficient that might be expected even if the indices were perfectly correlated with the proportion of species going extinct (Fig. S1). In the other cases, either the 95% confidence intervals of the observed correlations were greater than zero (Fig. 2), or the observed correlations were consistent with the range of values that might be expected (Fig S1).

The geometric mean abundance (*c*) of birds was estimated to be approximately 100,000 individuals per species. Assuming that the index based on the geometric mean is proportional to the extinction risk of species at the global scale, the reported decline in the LPI for birds from 1970 to 2009 of 13% (Baille et al. 2010) reflects a proportional increase in the probability of extinction of approximately 1% for values of *c* between 10,000 and 1,000,000 (Fig. 3). Smaller values of *c* imply larger changes in the risk of extinction for a given change in LPI, although the results are relatively insensitive to the choice of *c* (Fig. 3), and are primarily driven by the LPI values (Fig. S2).

**Figure 3.**
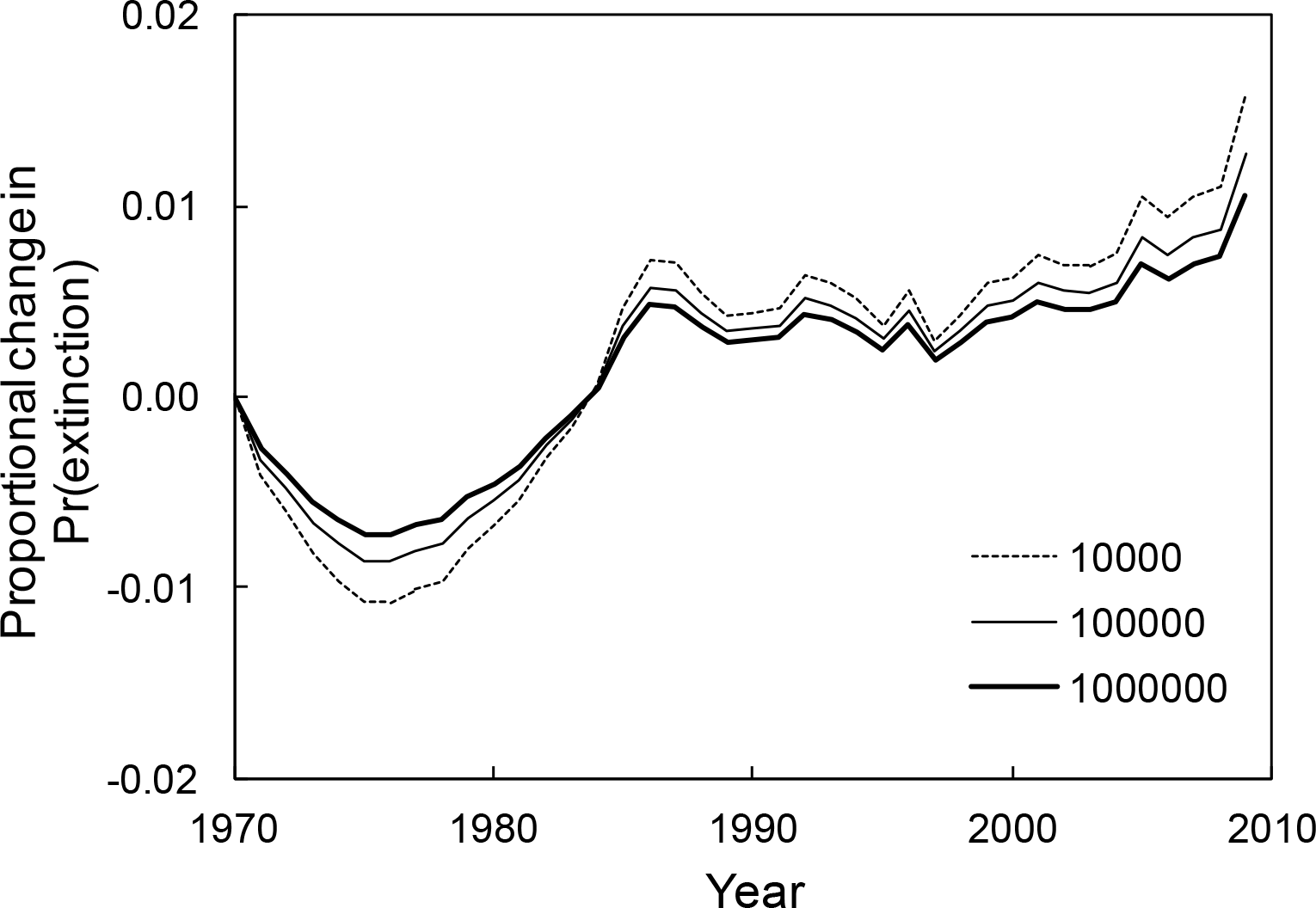
Proportional changes in the probability of extinction from levels in 1970 based on changes in the Living Planet Index for birds (Baille et al. 2010) assuming values for the geometric mean abundance in 1970 of 10,000, 100,000 or 1,000,000 individuals.

## Discussion

We derived indices that can be interpreted in terms of changes in extinction risk. By deriving the indices from theoretical population models, the merits of possible alternative indices can be assessed to determine which indices are best supported by data. Our analysis shows that the indices are positively correlated with the proportion of species going extinct in small patches, despite highly simplified assumptions used to build the indices.

In addition to the heuristic properties that Buckland et al. (2005) and van Strien et al. (2012) used to assess different indices of biodiversity, a good index should also be clearly related to particular management objectives. For example, we have shown that the geometric mean abundance of species, which has good heuristic properties (Buckland et al. 2005; van Strien et al. 2012), can be related to the proportion of species within an area that are likely to become extinct. This lends much greater support to this index as a biodiversity metric.

The geometric mean abundance of species is used increasingly, including in North American and European bird monitoring (Gregory & van Strien 2010; Butchart et al. 2010) and for planning fire management (Di Stefano et al. 2013). The LPI for reporting the state of species is the geometric mean abundance in each period, divided by the geometric mean abundance in the first time period (Loh et al. 2005; Collen et al. 2009). The LPI is based on the notion that changes in species abundance are important, but was not derived directly from ecological theory. We do not intend this as a particular criticism of the LPI, which has more support than some alternative indices, but we argue that ecological indices should have sound theoretical foundations. A theoretical foundation helps make the meaning and scope of the index clearer and more easily justified. For example, the derivation of the index based on the geometric mean implies that reductions in the LPI can be interpreted in terms of an increased average probability of extinction of the species. We estimate that the reduction of the global avian LPI of approximately 13% between 1970 and 2009 corresponds to approximately a 1% increase in the probability of extinction (Fig. 6). This is less than the increased risk of 7% implied by the Red List Index (RLI) of birds for the period 1988 to 2004 (Butchart et al. 2004), which is the only CBD index that is related directly to extinction. The larger increase in extinction risk implied by the RLI compared with *I*_g_ might be expected given the RLI’s focus on threatened species.

The indices based on the power mean (*I*_b_) and geometric mean (*I*_g_) have the same data requirements as those considered by Buckland et al. (2005). That is, they require information on the abundance of a suite of species at a particular point in time. The index that accounts for different trends among communities, (*I*_t_) requires additional information (the average trend of the species in the community). Such data will tend to be available for only a subset of species, and this subset is likely to be a biased sample of relevant species in a community. Any bias will be common to all indices, with the consequence that they might not broadly represent all possible species of interest.

Using a theoretical foundation to develop indices suggests ways in which the indices can be evaluated and improved, and assumptions underlying the indices are clear. The clear assumptions can be tested individually to determine whether they are violated in particular circumstances and the consequences of those errors. Further, the overall properties of an index can be assessed against data if it approximates an explicit quantity. In our case, we sought an index that would be linearly correlated with the proportion of species becoming extinct such that a change in the index would reflect a particular change in the proportion of species going extinct. The clear assumptions help highlight how the indices could be modified.

As an example of modification, trends in population size are likely to influence extinction risks. The index that incorporates trend (*I*_t_) shows how abundance and trend might be incorporated into a single index if the assumption of a consistent trend among communities is not supported. In the case of the experimental protozoan community, an assumption of an equal trend is clearly not supported. Of the four protozoan species, one went extinct in all 40 experimental replicates, and one persisted in all replicates. Thus, the proportion of species in each community that went extinct was influenced substantially by the identity of the species, which had different trends not just different population sizes.

Biodiversity indices, such as those developed here, will be sensitive to the choice of species that are included. For example, species included in the LPI calculations are not a random sample of all possible species, with biases likely. Unless the scheme used to select the sample of species used in the index is considered carefully, it will be unclear how the selected species will represent the broader suite of biodiversity.

Factors other than those included in the indices are likely to influence extinction. The Lepidoptera species will be differentially susceptible to apparent local extinction because of different dispersal and abilities to persist outside the focal habitat patches. Other species will occur only ephemerally in the patches, reducing the influence of abundance on local extinction. However, the results were qualitatively identical when analysing only strict grassland specialists, so we reported only the results for the larger collection of species.

Our indices were based on models of exponential decline of single populations, thereby ignoring spatial aspects and density-dependence. Other indices based on metapopulation dynamics, for example, could be developed to account for spatial effects. Indeed, metapopulation capacity, which is derived from colonisation and extinction dynamics of habitat patches (Day & Possingham 1995; Hanski & Ovaskainen 2000), can be viewed as an index of metapopulation persistence (Moilanen & Nieminen 2002). Density-dependence might be less important for populations that are declining deterministically, although accounting for non-exponential decline might be important because temporal patterns of decline influence risk (Di Fonzo et al. 2013).

Imprecise estimation of abundance (particularly in the woodland case study), some residual uncertainty about the local extinction of species due to imperfect detection, and the false assumption of equivalent dynamics of all species would all weaken the correlation between the indices and the observed extinction rate. Despite this, the predicted and observed extinction risks were correlated (Figs 1–3). This implies that using the indices to aggregate data across species is reasonable. However, further tests of the indices to predict local extinction would be valuable, as would evaluating extinction risk over regions larger than just single patches (e.g., based on spatial population dynamics).

The index based on the power mean is sensitive to the choice of the parameter *b*, and estimating it via estimates of the standard deviation of the population growth rate (*σ*) might be difficult. Thus, the indices based on the geometric mean (*I*_g_ and *I*_t_) might be more appealing because a freely-varying parameter does not require estimation.

Further, extinctions might be dominated by deterministic declines rather than random fluctuation around a zero mean growth rate. If true, the indices based on the geometric mean might be preferred over that based on the power mean.

The SAFE index (Clement et al 2011; see also Akçakaya et al. 2011; Beissinger et al. 2011; McCarthy et al. 2011) is essentially equal to the logarithm of population size. Our analysis shows, therefore, that the SAFE index will be proportional to the expected time to quasi-extinction (time to reaching a given threshold). But it also shows that the SAFE index will be comparable among species as a measure of threat only if trends in population size of those species are similar. Where trends differ among species, an index based on–ln[*x*(0)]*/*ln[*λ*] is likely to better reflect threat. Further, prioritization of management, which apparently motivated the SAFE index, should not be based on extinction risk, but on the ability to change risks (McCarthy et al. 2011). This might be assessed, for example, by the relative cost of changing *x*(0) or *λ* and their influence on–ln[*λ*]/ln[*x*(0)] (Baxter et al. 2006).

An index developed without theory does not mean it will have poor properties. As we have seen, the geometric mean was developed without theory but appears to have useful properties (Buckland et al. 2005; van Strien et al. 2012). The demonstrated relationship to extinction risk lends further support to the geometric mean. Our analysis also indicates how the geometric mean might incorporate population trends. We suggest that biodiversity indices should be developed more frequently from theoretical foundations to provide more explicit links between the index, the data underlying the index, and the meaning of changes in the index. Such indices will inevitably exclude factors that might be important; this is a feature of any model.

However, stronger theoretical foundations for biodiversity indices would clarify the features that are considered and those that are ignored, and would allow the indices to be more easily evaluated and improved.

## Acknowledgments

This research was supported by an Australian Research Council (ARC) Future Fellowship to M.A.M. (FT100100923), and the ARC Centre of Excellence for Environmental Decisions. We thank J. Baumgartner for help with collating the data on bird abundances, B. Collen for providing values for the Living Planet Index of birds, and J. Moore, R. Gregory, R. Camp and two anonymous reviewers for comments on a draft manuscript.

## Supporting Information

Further details on the datasets used (Appendix S1), details on calculating the expected correlation (Appendix S2), and Figs S1 and S2 are available online. The authors are solely responsible for the content and functionality of these materials. Queries (other than absence of the material) should be directed to the corresponding author.

